# The role of type IV pilus in the interaction of *Neisseria gonorrhoeae* with a corneal epithelium tissue model

**DOI:** 10.64898/2025.12.22.695979

**Authors:** Lina Kafuri, Nicola Knetzger, Christian Lotz, Vignesh Parasuraman, Thomas Rudel, Vera Kozjak-Pavlovic

## Abstract

*Neisseria gonorrhoeae*, the causative agent of the sexually transmitted disease gonorrhea, can infect intact corneal epithelium, leading to inflammation, cornea perforation, and blindness. Models for studying this type of gonococcal infection are few and limited. We have tested cornea models based on an immortalized hTCEpi corneal epithelium cell line for infection research using four derivatives of the *N. gonorrhoeae* MS11 strain that differ in pilus expression. We show that bacterial adherence depends on the functional type IV pilus in the early stages of infection, and that the formation of bacterial microcolonies and biofilms on the model surface leads to tissue destruction. The infection induces a specific cytokine response characterized by an increase in the secretion of IL-8 and TNF-α, but not of IL-6. The testing of trifluoperazine, a drug that induces pilus retraction, on infected corneal tissue models showed that the drug strongly diminished the number of adherent gonococci only when applied simultaneously with bacteria, and not when bacteria were allowed to form microcolonies on the tissue surface. Our work describes, for the first time, hTCEpi-based corneal epithelium tissue models as a useful tool for investigating *N. gonorrhoeae* infection, with potential for application in high-throughput studies and drug screening.

## Introduction

*Neisseria gonorrhoeae* is a gram-negative diplococcus that causes the sexually transmitted disease gonorrhea. The infection in women is usually asymptomatic, and the risk of transmission to the newborn during birth is considerably high for undiagnosed pregnant women, which is often the case in countries where the screening for *N. gonorrhoeae* is lacking (1). Gonococcus (GC) can adhere and colonize the intact corneal epithelium, and is the main cause of *ophthalmia neonatorum*, an aggressive eye infection characterized by the inflammation of the eyelids and the presence of a yellow discharge, which can lead to cornea ulceration, perforation, and blindness (2). Adults are also susceptible to gonococcal eye infections. In recent years, the number of reported cases of eye infections caused by multi-antibiotic-resistant GC has increased due to misdiagnosis of the causing pathogen, delaying proper treatment and leading to permanent cornea injuries (1).

*N. gonorrhoeae* uses different virulence factors to adhere to epithelial cells, including the opacity proteins (Opa) and the type IV pilus. This complex protein machinery consists of many subunits, which together make dynamic filaments with the possibility of retraction and extrusion (3). The main subunit, pilin or PilE, has a high intra-strain structural variation dependent on the homologous recombination of the large family of PilS silent genes, mediated by the recombinase A (RecA) (4). This high variation not only allows *N. gonorrhoeae* to escape the immune system but also affects how they adhere to epithelial cells. However, PilE is not the only determining factor of the adherence behavior of GC (5). PilC, encoded by two different genes, pilC1 and pilC2, is located on the tip of the pilus where it acts as an adhesin, thus allowing the binding of the bacterium not only to the target tissue (6) but also to other bacteria, which facilitates the formation of microcolonies.

*N. gonorrhoeae* infection research, especially in the genitourinary tract, has been carried out using various models, including humanized hormone-treated mice, *in vitro* models using 2D cell monolayers, and 3D tissue models and organoids. This has allowed a better understanding of the key factors in gonococcal adherence and invasion (7). Previous infection studies focusing on corneal tissue mostly used human corneal explants (8) or the culture of human corneal epithelial cells on a bovine cornea-derived scaffold (1). Of the different human corneal tissue models established so far, most have been made as an alternative to the Draize irritation test (9), but not specifically for the infection studies. The accessibility and high variability of the patient-derived primary human corneal epithelial cells can make these models difficult to make and standardize. As an alternative, different human corneal epithelial cell lines can be used, such as a telomerase-immortalized human corneal epithelial cell line (hTCEpi), which can form stratified epithelium expressing corneal differentiation markers like ZO-1 and cytokeratin 3 when cultured under air-lift conditions (10).

In this work, we used an hTCEpi-based corneal model to study GC infection. We assessed the role of the type IV pilus, using derivatives of the MS11 strain with differences in pilus expression and adherence: MS11 F3 Pil- (Pil-, Opa+, PilC+, recA+), MS11 F3 Pil+ (Pil+, Opa+, PilC+, recA+), MS11 N159 (Pil+, Opa+, PilC+, recA-), and MS11 N191 (Pil+, Opa+, PilC-, recA+). We analyzed the effect of GC infection on tissue permeability, cytotoxicity, and the production of key cytokines, as well as demonstrated the importance of microcolony formation for the damage of the corneal tissue, which is a symptom often observed during GC infection in patients. Lastly, we tested the effect of trifluoperazine, a drug that can induce pilus retraction in *Neisseria meningitidis*. We assessed the pilus retraction effect of the drug in terms of bacterial adherence and IL-8 secretion. We also analyzed the bactericidal activity of trifluoperazine and the induction of GC microcolony disaggregation. In sum, the cornea model used in this study is a suitable and reproducible tool not only for studying basic aspects of the GC eye infection but also for the testing of antimicrobials.

## Materials and methods

### Generation of corneal epithelium tissue models

Human corneal epithelium tissue models were derived from telomerase-immortalized human corneal epithelial cell line (hTCEpi) (Evercyte, Vienna, Austria, #CHT-045-0237) (10). These cells were maintained in E1 medium (EpiLife medium) (Thermo Fisher Scientific, Massachusetts, USA, #MEPI500CA) supplemented with 1 % human keratinocyte growth supplement (HKGS) (Thermo Fisher Scientific, Massachusetts, USA, #S0015) and 1 % penicillin/streptomycin (P/S) (Sigma-Aldrich, Darmstadt, Germany, #P4333). Cells were seeded into culture flasks at a density of 4–6 cells/cm² and grown to 80–90 % confluence under standard conditions (37 °C, 5 % CO₂, 95 % humidity). The medium was exchanged every 2–3 days.

For passaging, cells were detached with trypsin and centrifuged at 200xg for 5 min. The pellet was resuspended in E2 medium (EpiLife medium supplemented with 1 % HKGS, 1 % P/S, and 0.48 % 300 mM CaCl₂).

Human corneal epithelium models were cultured in BRAND cell culture inserts (BRAND, Wertheim, Germany, #782700) with polycarbonate membranes (0.4 µm pore size and 0.59 cm² growth area). For seeding, 3x10LJ cells in 300 µl E2 medium were added to the apical compartment of each insert and incubated for 2 h under standard conditions (37 °C, 95 % humidity, 5 % CO₂). Subsequently, 1.4 ml E2 medium was added to the basolateral compartment. After 24 h of submerged culture, the medium on the apical side of the inserts was removed, while the medium on the basolateral side was replaced with 1.4 ml E3 medium (EpiLife medium supplemented with 1 % HKGS, 1 % penicillin/streptomycin, 0.48 % CaCl₂, 73 µg/ml ascorbyl-2-phosphate and 10 ng/ml KGF) to initiate air–liquid interface culture. The medium was exchanged every 2–3 days thereafter.

### Histology

Corneal epithelium models were fixed for 2 h with 4 % paraformaldehyde (Morphisto, Offenbach am Main, Germany, #11762.01000) and, after washing three times with 1xPBS, they were disassembled using an 8 mm biopsy puncher. After paraffin embedding, the samples were sectioned into 7 µm-thick slides. After a deparaffinization process including xylene and rehydration with a gradient of different concentrated solutions of ethanol, hematoxylin and eosin staining was done according to standard protocols (11). The stained tissue slides were analyzed with a light microscope (Leica Microsystems, Wetzlar, Germany).

### Neisseria gonorrhoeae strains

All the *N. gonorrhoeae* strains used in this study are MS11 derivatives, and their whole genomes were sequenced using Oxford Nanopore (Microsynth, Balgach, Switzerland). The sequences described in this publication are available at NCBI’s BioProject database under the accession number PRJNA1356842.

*N. gonorrhoeae* F3 Pil- (Pil^-^, PilC^+^, Opa^+^, recA^+^), F3 Pil+ (Pil^+^, PilC^+^, Opa^+^, recA^+^), N159 (Pil^+^, PilC^+^, Opa^+^, recA^-^), and N191 (Pil^+^, PilC2, Opa^+^ , recA^+^) were grown for 24 h at 37 °C and 5 % CO_2_ on plates with Oxoid GC agar base (Thermo Fischer Scientific, Massachusetts, USA, #CM0367) supplemented with 1 % (v/v) vitamin mix. The day after, approximately 10 single colonies were selected, re-streaked onto a new plate, and grown for 16 h before being used for infection.

### Infection of corneal epithelium models

Human corneal epithelium models were infected with the different *N. gonorrhoeae* strains on day 7 of the development of the models. The bacteria from the plate were taken with a sterile cotton swab and resuspended in 1 ml of DMEM F12 medium (Gibco, Thermo Fisher Scientific, Massachusetts, USA, #31331093). The OD_550_ was measured to calculate the bacterial number, and, taking into account the number of epithelial cells, the models were infected from the apical side for 2 h at an MOI of 20 in DMEM F12 medium. The models were then washed three times with 1xDPBS and incubated under airlift conditions until further evaluation at different time points (2 h, 24 h, and 72 h).

### Immunoblotting

PilE (pilin) expression was analyzed by sodium dodecyl sulfate polyacrylamide gel electrophoresis (SDS-PAGE) and western blot. Briefly, the samples were dissolved in Laemmli buffer (62.5 mM Tris pH 6.8, 2 % SDS, 10 % glycerol, 5 % β-mercaptoethanol, and 0.002 % Bromophenol Blue) and separated on a 4-12 % NuPAGE Bis-Tris Precast gel (Thermo Fisher Scientific, Massachusetts, USA, # NP0322BOX). The separated proteins were then transferred onto a polyvinylidene fluoride (PVDF) membrane, which was blocked for 1 h with 5 % skimmed milk in PBS, followed by the overnight incubation with the pilin SM1 mouse monoclonal antibody (a kind gift from Magdalene Yh So, University of Arizona), Omp85 (custom-made in rabbits against the full-length protein; Davids Biotechnology, Regensburg, Germany), or opa (in-house made in rabbits) at 4 °C. For detection, the membrane was incubated with an HRP-coupled secondary antibody (Biozol, Eching, Germany), and the images were processed with FIJI (12).

### Barrier integrity of corneal epithelium models

To test the tissue integrity of the human cornea models upon infection, we measured the permeability to 4 kDa fluorescein isothiocyanate (FITC)-Dextran (Sigma Aldrich, Darmstadt, Germany, #46944). For this, 0.25 g/ml FITC-Dextran was dissolved in Dulbecco’s Modified Eagle Medium (DMEM) (Sigma Aldrich, Darmstadt, Germany, #D6429) and sterile filtered. The medium on the basolateral side of the insert was removed and replaced with 1 ml of DMEM, while 300 µl of the FITC-dextran solution was added to the apical side. The models were incubated for 30 min (37 °C and 5 % CO_2_), and then 200 µl from the basolateral side was transferred to a black clear-bottom 96-well plate (Corning, New York, USA, #3603). The fluorescence was analyzed using a TECAN reader (490 nm absorption and 525 nm emission), and the results were normalized to the values measured from an empty cell culture insert.

### Cytotoxicity

Cytotoxicity was assessed by measuring the lactate dehydrogenase (LDH) activity in the cell supernatant using the Cytotoxicity Detection KitPlus (Roche, Basel, Switzerland, #4744926001). The assay was performed according to the manufacturer’s instructions, using the medium from the basolateral side of the models.

### Bacterial adherence

After the different time points, the medium was removed from the basolateral side of the cell culture insert, and using an 8 mm biopsy puncher, the membrane was carefully removed and placed into an Eppendorf tube containing 300 µl of 1 % (w/v) saponin solution (Sigma Aldrich, Darmstadt, Germany, #47036), and the tubes were vortexed for 1 min. After 30 min of incubation, serial dilutions were made and plated on GC agar plates. The CFU/ml was determined after 48 h of incubation at 37 °C with 5 % CO_2_.

### Cytokine measurements

72 h after infection, the medium was collected from the basolateral side of the cell culture insert and analyzed using a personalized Legendplex cytokine panel assay kit (BioLegend, San Diego, USA) to detect and measure the concentration of IL-1β, TNF-α, IL-6, IL-8, IL-10, and MCP-1. The assay was performed according to the manufacturer’s instructions, and the results were analyzed using the BioLegend software.

For the drug treatment experiments, the concentration of secreted IL-8 was measured in the medium collected from the basolateral compartment of the models infected with *N. gonorrhoeae* MS11 N159 and simultaneously treated with 40 µM trifluoperazine (SimTr). For this, the Human IL-8 DuoSet ELISA kit (R&D Systems, Minneapolis, USA, #DY208) was used, and the analysis performed according to the manufacturer’s instructions.

### Immunofluorescence staining and imaging

At different time points, the medium was removed from the basolateral side of the cell culture inserts, and the infected corneal epithelium models were fixed with 4 % paraformaldehyde (Morphisto, #11462.01000) for 2 h at room temperature, followed by washing 3 times with PBS and blocking with 3 % (w/v) bovine serum albumin (BSA) (Carl Roth, Karlsruhe, Germany, #8076.3) and 0.01 % Triton X (Carl Roth, Karlsruhe, Germany, #3051.4) in PBS for 1 h. The models were then incubated with primary antibody against *N. gonorrhoeae* (United States Biological, Salem, USA, #N0600-02) at 4 °C overnight. This was followed by incubation with a fluorophore-coupled secondary antibody (Alexa Fluor 488, Thermo Fisher Scientific, Massachusetts, USA, #A-11008), Phalloidin-Alexa Fluor 555 (Invitrogen/ Thermo Fisher Scientific, Massachusetts, USA, #A34055), and DAPI (Sigma Aldrich, Darmstadt, Germany, #D9542). The samples were mounted using Mowiol® (Carl Roth, Karlsruhe, Germany, #0713.2). For imaging, Z-stacks were made using a Leica STELLARIS 5 confocal imaging platform (Leica Microsystems, Wetzlar, Germany), and the images were processed using FIJI (12).

### Scanning Electron Microscopy (SEM)

After 72 hours of infection, the membranes of the cell culture inserts were carefully removed with an 8 mm biopsy puncher and were fixed with 6.5 % glutaraldehyde at 4 °C overnight. Subsequent sample preparation was done as previously described (13) and imaging was performed using a JEOL JSM-7500F microscope.

### Drug assays

We applied two different approaches to study the effects of trifluoperazine on pilus retraction. For the first approach, post infection treatment (PostTr), the corneal epithelium models were infected with *N. gonorrhoeae* MS11 N159 for 1 h, as described previously in this study. The models were washed 3 times with DPBS to remove the non-adhered bacteria and were then treated with 40 µM trifluoperazine (Sigma Aldrich, Darmstadt, Germany, #T8516) for 1 h. After this time, the models were washed 3 times with DPBS and were left under the airlift condition for another 72 h. For the second approach, the simultaneous addition of bacteria and drug (SimTr), the models were infected with *N. gonorrhoeae* MS11 N159 as described and treated at the same time with 40 µM trifluoperazine. 1 h later, the models were washed 3 times with DPBS and left under the airlift condition for further 72 h. Bacterial adherence was then measured as previously described in this study.

### Bacterial growth curves

The *N. gonorrhoeae* strains were grown for 24 h on GC agar plates. After this time, they were re-streaked onto a fresh GC agar plate for 16 h before the experiment. The bacteria were then taken up with a sterile cotton swab and diluted in 1 ml PPM+ medium (15 g/l proteose peptone (Becton Dickinson, New Jersey, USA, #211693), 5 g/l sodium chloride (VWR, Radnor, USA, #27810.364), 0.5 g/l soluble starch (Sigma-Aldrich, Darmstadt, Germany, #33615), 1 g/l potassium dihydrogen phosphate (Carl Roth, Karlsruhe, Germany, #3904.1), and 4 g/l dipotassium hydrogen phosphate (Carl Roth, Karlsruhe, Germany, #P749.3), supplemented with 1 % (v/v) vitamin mix and 0.5 % (v/v) sodium hydrogen carbonate (VWR, Radnor, USA, #27775.293). The OD_550_ was measured and adjusted to 0.3 in 20 mL PPM+ medium and grown at 37 °C with constant shaking (130 rpm), until they reached an OD_550_ of at least 0.4 (exponential phase). Then they were diluted to an OD_550_ of 0.1, with 40 µM trifluoperazine added to the corresponding cultures. The OD_550_ was measured every hour for 5 h.

### Bactericidal assay

To test the bactericidal effect of different concentrations of trifluoperazine in the two approaches (PostTr and SimTr), *N. gonorrhoeae* MS11 N159 was grown on GC agar plates for 24 h and then re-streaked onto a fresh plate, where it grew for approximately 16 h. The bacteria were then taken up with a sterile cotton swab, resuspended in 1 ml DMEM F12 medium, and the OD_550_ was measured. The same volume needed to infect the corneal epithelium models with an MOI 20 was calculated, and the solutions were prepared accordingly, with the different concentrations of trifluoperazine (10, 20, 30, 40, and 50 µM) and gentamicin (150 µg/ml) (Gibco, Thermo Fisher Scientific, Massachusetts, USA, # 15710-049). 100 µl of each solution was added to a flat-bottom 96-well plate.

For the first approach (PostTr), the plate containing just bacteria was incubated for 1 h at 37 °C with 5 % CO_2_; after this time, trifluoperazine and gentamicin were added to the corresponding wells, and the bacteria were treated for 1 h.

For the second approach (SimTr), the plate containing the bacterial suspension with different concentrations of trifluoperazine and gentamicin was incubated for 1 h at 37°C with 5 % CO_2_.

After incubation, serial dilutions were made and plated onto GC agar plates. Colonies were counted after 48 h to calculate the CFU/ml.

### Aggregation assay

To analyze the effect of trifluoperazine on the disaggregation of microcolonies, *N. gonorrhoeae* MS11 N159 was grown for 24 h on a GC agar plate, re-streaked onto a fresh GC agar plate, and grown for approximately 16 h. The bacteria were collected with a sterile cotton swab, diluted in 1 ml PPM+ medium, and the OD_550_ was measured. The OD was adjusted to 0.3 in 15 ml PPM+ medium, and bacteria were grown at 37 °C with constant shaking (130 rpm) until they reached an OD_550_ of approximately 0.4. The bacterial culture was diluted with the DMEM F12 medium to an OD of 0.1. 500 µl of this suspension was distributed into a 24-well plate and incubated (37 °C, 5 % CO_2_) for 2 h. After this time, the bacteria were treated with increasing concentrations of trifluoperazine (10, 20, 30, 40, and 50 µM) and with gentamicin (150 µg/ml) for 1 h. Before and after the treatment, three images per well were taken using a Leica DMIRB microscope (Leica Microsystems, Wetzlar, Germany), and the area of the microcolonies was calculated using FIJI (12).

### Statistical methods

Statistical analyses were performed on at least three independent biological replicates using 2-way ANOVA and Tukey’s multiple comparison tests, with the help of the GraphPad Prism software version 10.4.2.

## Results

### Type IV pilus plays a crucial role in the adherence of *N. gonorrhoeae* to the corneal epithelium

To study the role of the type IV pilus of *N. gonorrhoeae* in the adherence to the intact corneal epithelium, we used a multilayer model made of immortalized human derived corneal epithelial cell lines (hTCEpi) seeded onto polycarbonate membrane cell culture inserts. These models show corneal differentiation traits, and their architecture resembles the native tissue (Fig. 1A) (10). We allowed bacteria to interact with the tissue for 2 h before washing and removing the ones that did not adhere. Further analyses of the models were then performed at different infection times, starting directly after the wash (2 h time point) and finishing 72 h later (Fig. 1B).

**Figure 1.**
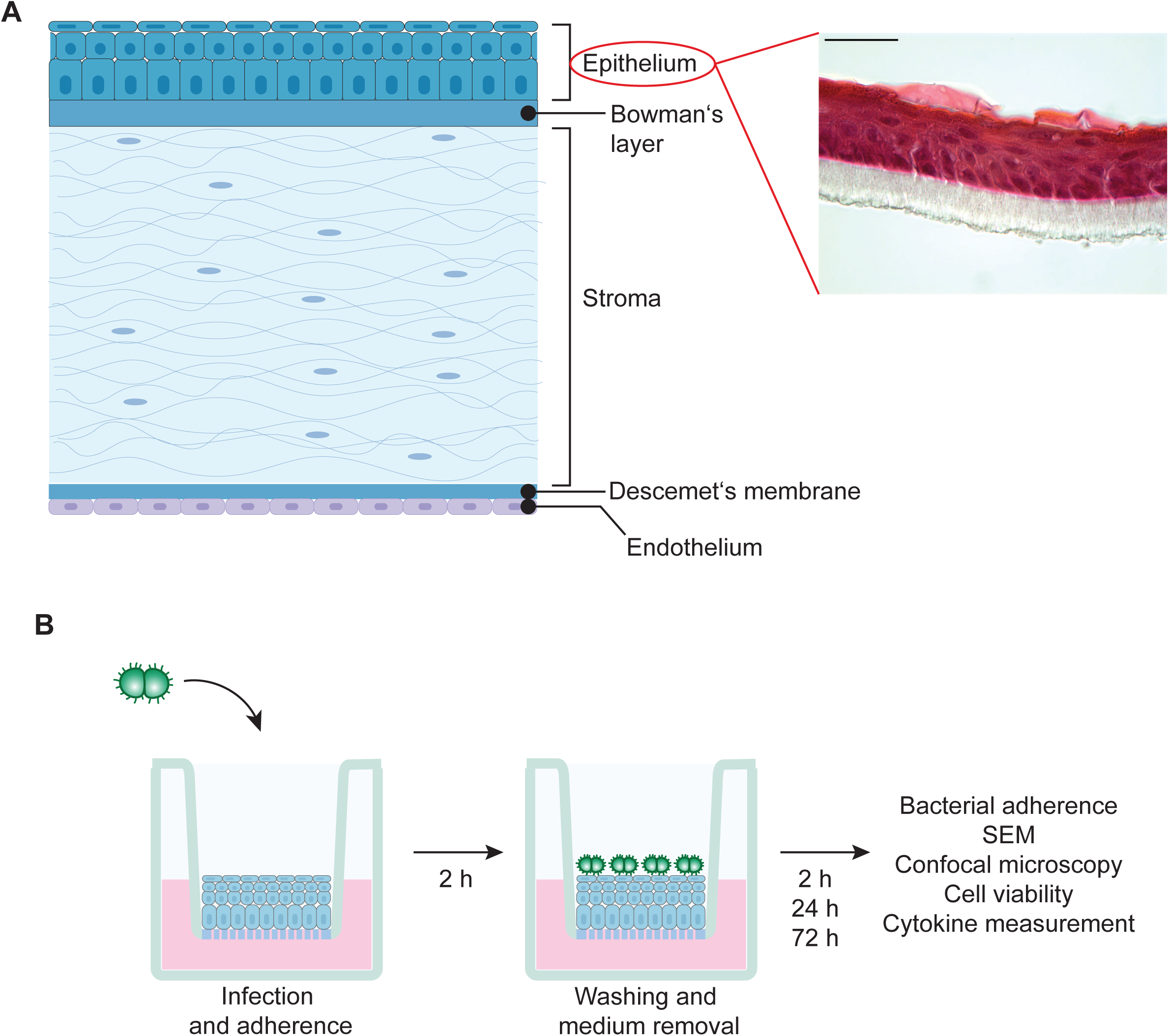
Schematic representation of the corneal epithelium structure and study design. (A) The scheme of the cornea structure (on the left side) in comparison to the corneal epithelium model used in this study. The corneal tissue model was fixed, embedded in paraffin, and sectioned into 7 µm-thick sections, stained using hematoxylin and eosin (scale bar 50 µm). (B) Corneal tissue models were infected from the apical side with the different *N. gonorrhoeae* MS11 derivative strains for 2 h, followed by washing with PBS and further incubation under airlift conditions. Different parameters were analyzed at different time points.

We infected the models with four different *N. gonorrhoeae* strains derived from MS11, which exhibit different pilus phenotypes (Fig. 2A). To assess these differences, we first performed whole genome sequencing of each strain and analyzed the regions encoding for important proteins like pilE, pilC, opacity-associated (opa) proteins, and recA (Table 1). We observed that one of the two pilE genes in the F3 Pil- strain was not present and the other one was partially deleted, resulting in no expression of pilin, which was confirmed by western blot of bacteria before and after 2 h of infection (Fig. 2A, Supp. Fig. 1). This gene was intact in F3 Pil+, N159, and N191; however, there was a noticeably higher expression of the pilus in N159 (Fig. 2A). In addition, recA gene in N159 is inactivated due to a large insertion, which drastically reduces phase variation of the pilus. N191, although possessing an intact pilE gene, has mutations in both pilC genes, which strongly diminished the adherence of these bacteria (Table 1).

**Figure 2.**
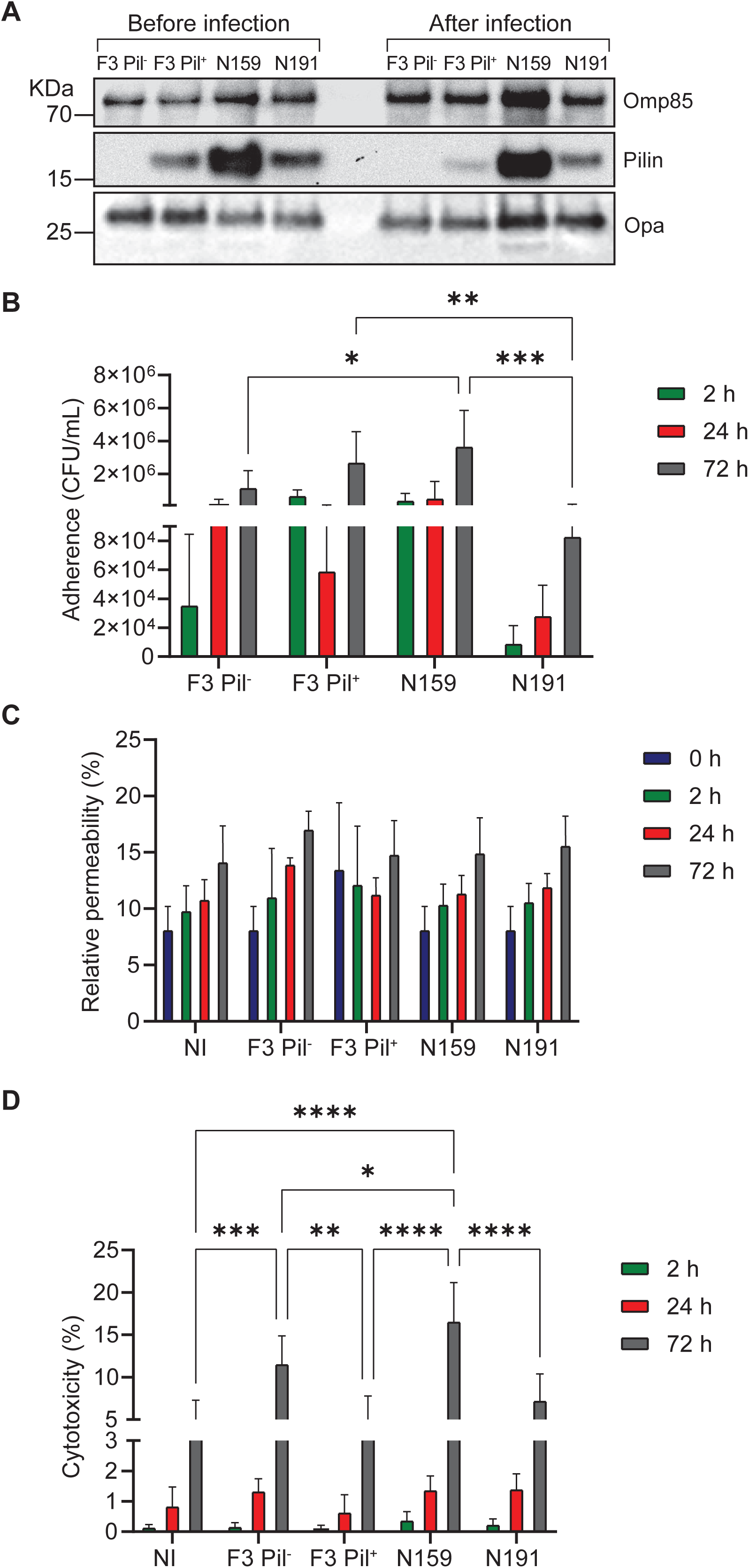
Impact of pilus expression on bacterial adherence, corneal tissue model barrier integrity, and cytotoxicity. (A) The expression of Pilin in different *N. gonorrhoeae* MS11 derivative strains was analyzed by immunoblotting before and after 2 h of infection using antibodies against Pilin, Omp85 (Outer membrane protein assembly factor BamA) and opa. (B) To determine the number of adherent bacteria, corneal tissue models were solubilized with 1 % saponin for 30 min. Serial dilutions were plated on GC agar to calculate the CFU/ml. (C) The corneal epithelium barrier integrity was assessed before (0 h) and at different time points of infection by measuring the diffusion of a FITC-dextran solution from the apical to the basolateral compartment of the models. The values were normalized to the values of an empty cell insert. (D) Cytotoxicity during different time points of infection was measured as the percentage of lactate dehydrogenase LDH released to the medium collected from the basolateral side of the corneal epithelium models. All graphs represent mean values ± SD from three independent replicates. Statistical analysis was done using 2-way ANOVA and Tukey’s multiple comparison tests. * - p≤0.05, ** - p≤0.01, *** - p≤0.001, and **** - p≤0.0001.

**Table 1.**
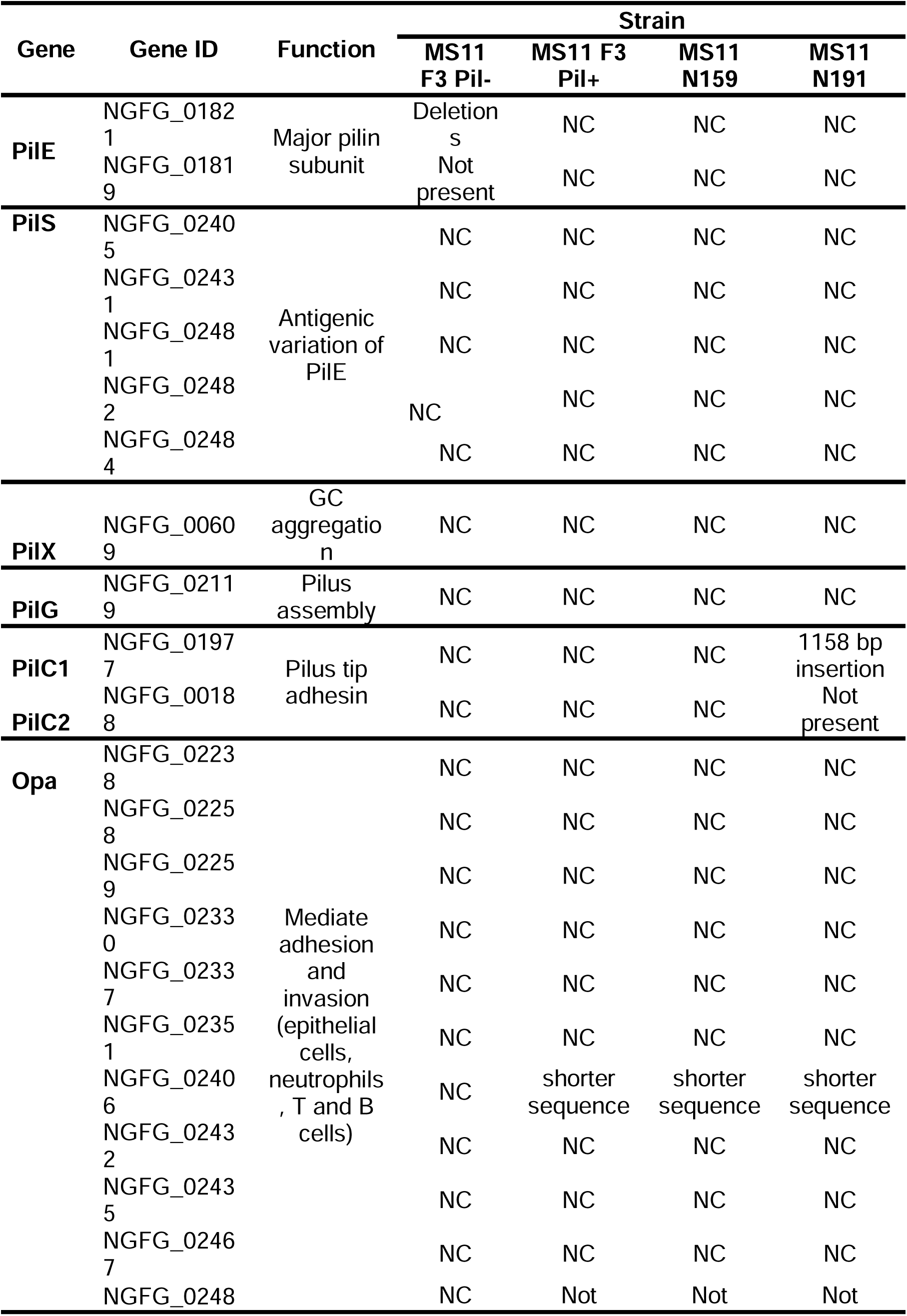
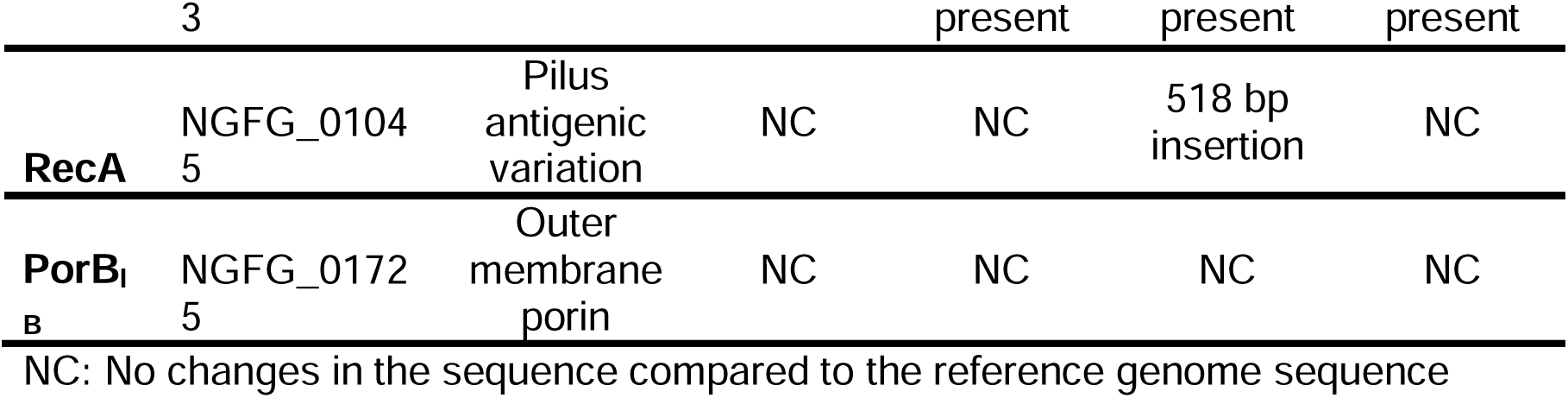
Comparison of relevant genes involved in bacterial adherence and invasion between the different *N. gonorrhoeae* MS11 derivative strains and the reference genome of *N. gonorrhoeae* MS11 (NCBI, CP003909) according to the information obtained by whole genome sequencing.

We assessed bacterial adherence, cytotoxicity, and tissue permeability during infection. N159 strain showed the highest adherence at all time points (2 h, 24 h, and 72 h). The adherence capacity of F3 Pil+ was the second highest (Fig. 2B). This strain resembles the wildtype MS11 genotype and contains the intact recA gene, so the pilus undergoes phase-variation at a much higher rate compared to N159; this was reflected in the fact that the pilus expression in the bacteria collected after 2 h of infection was reduced (Fig. 2A), indicating that nonadherent bacteria were less piliated. F3 Pil- strain showed a somewhat reduced adherence capacity, as expected, due to the lack of PilE expression (Fig. 2A, B), especially at 2 h. Lastly, the N191 strain showed the lowest adherence to the corneal epithelium, emphasizing the importance of PilC in the adhesion to the target tissue (6) (Fig. 2B).

The mature models at the time of infection have a relative permeability of 8 %, as measured by the FITC-dextran assay. The permeability increased over time to up to 15 % in both uninfected and infected models, indicating that this was probably caused by the aging of the tissue models and not by infection (Fig. 2C).

The cytotoxicity upon infection was measured in relation to the concentration of lactate dehydrogenase (LDH) released into the medium of the basolateral compartment. This is connected to cell lysis, usually associated with apoptosis or necrosis. We observed an increase in cytotoxicity with time, showing the highest percentage in the models infected with the N159 strain for 72 h (17 %). Interestingly, the F3 Pil- strain showed the second highest cytotoxicity, followed by F3 Pil+ and N191, for which the values were comparable to those obtained for the non-infected models (Fig. 2D).

In conclusion, adherence and cytotoxicity mostly correspond to the ability of bacteria to adhere to the corneal tissue models with the help of the pilus. Relatively high adherence and cytotoxicity of F3 Pil- strain indicate, however, that bacteria can use other mechanisms besides pili to efficiently adhere to the corneal tissue.

### *N. gonorrhoeae* forms microcolonies on the surface of corneal tissue models, depending on the presence of the pilus

Another important role of the type IV pilus in *N. gonorrhoeae* is the formation of microcolonies. They allow GC to survive longer in the target tissue in the presence of some antibiotics, due to their limited diffusion within the microcolony (14).

We observed large microcolonies already after 2 h of infection for the piliated strains F3 Pil+ and N159. These microcolonies were distributed over the whole surface of the corneal epithelium models. In contrast, F3 Pil- and N191 bacteria were present on the models as single bacteria or in much smaller aggregates, visible only after 72 h of infection (Fig. 3A).

**Figure 3.**
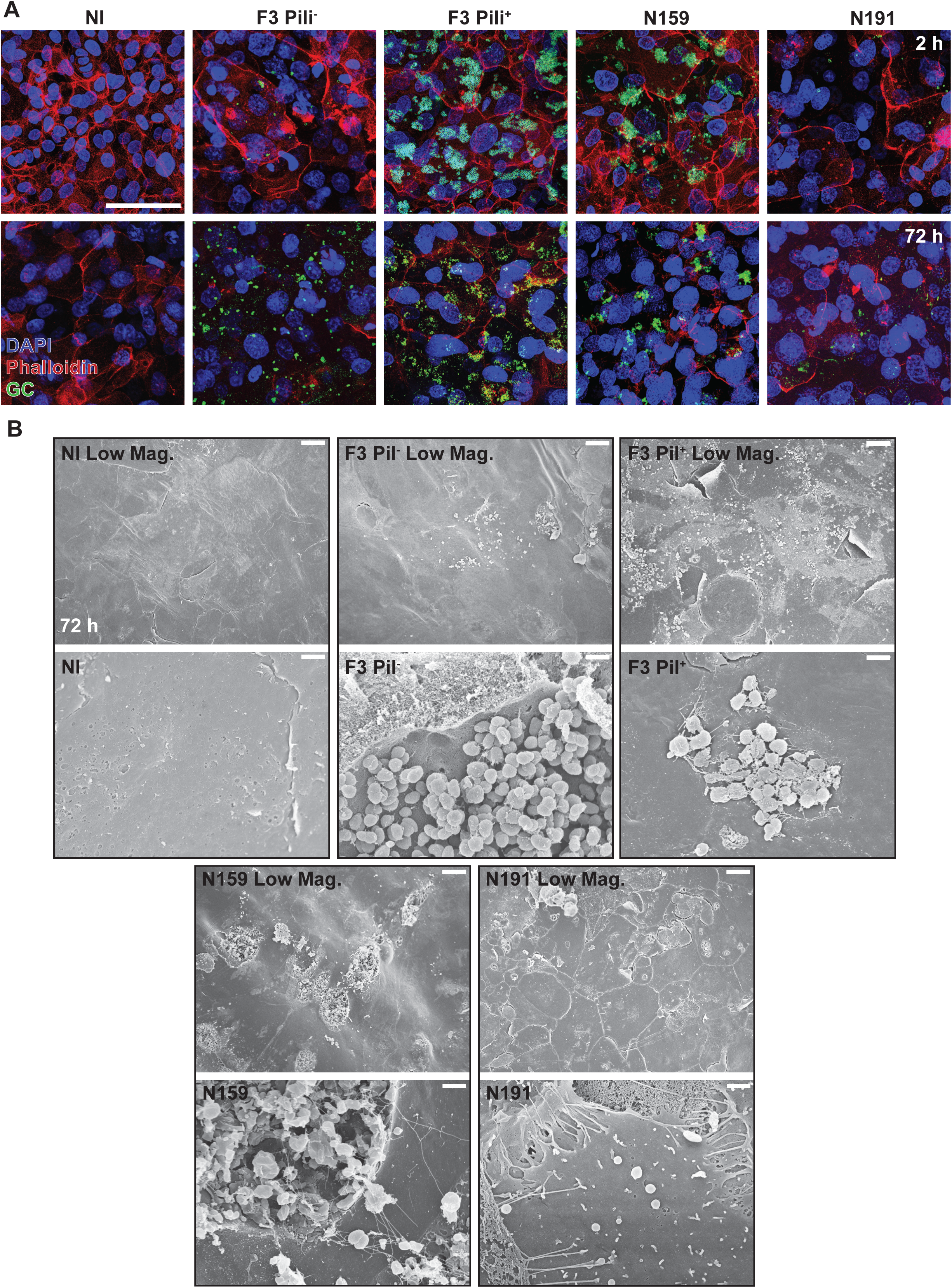
Pilus-expressing *N. gonorrhoeae* MS11 derivative strains form microcolonies on the surface of corneal tissue models. (A) The models were fixed 2 h and 72 h post-infection and analyzed by immunofluorescence and confocal microscopy, using DAPI (blue), Phalloidin-Alexa Fluor 555 (red), and the primary antibody against *N. gonorrhoeae*/Alexa Fluor 488-coupled secondary antibody (green). The images were made using the Leica Stellaris 5 confocal imaging platform. Scale bar is 50 µm. (B) Corneal tissue models infected with different *N. gonorrhoeae* MS11 derivative strains were fixed after 72 h of infection and analyzed by scanning electron microscopy (SEM). Scale bars are 10 µm for low magnification (Low Mag.) and 1 µm for other images.

Scanning electron microscopy (SEM) of the infected tissue models 72 h post infection revealed that F3 Pil+ and N159 microcolonies led to tissue damage, with pits forming underneath bacterial aggregates attached and connected through long filaments, most likely representing pili. Such filaments were absent in F3 Pil-, though we observed occasional accumulations of these bacteria resembling a biofilm. N191 were hard to detect in SEM images and were found only as single diplococci attached to the cell surface. Interestingly, pili filaments of N159 appeared much longer and more abundant than those of F3 Pil+ (Fig. 3B).

Microscopy shows, therefore, that only piliated bacteria form large microcolonies, which lead to tissue damage. Consequently, the high cytotoxicity observed for the N159 strain corresponds to the most pronounced tissue damage visible on the SEM images.

### Corneal epithelial cells produce different cytokines and chemokines upon infection with *N. gonorrhoeae*

Previous studies have shown that bacteria induce the secretion of different cytokines by the mucosal epithelial cells, which are the first responders to infection. The secreted cytokines regulate other cell types, recruiting immune cells to the site of infection. The degree of this response is dependent on the cell type, as well as on the pathogen and its virulence factors (15).

We evaluated the secretion of different cytokines and chemokines after 72 h of infection with the four strains we used. Interleukin (IL)-8 was the most secreted cytokine, followed by Monocyte Chemoattractant Protein (MCP)-1, IL-1β, and Tumor Necrosis Factor (TNF)-α, whereas IL-6 and IL-10 were not detectable, or present in very low amounts (below 3 pg/ml) (Fig. 4A). The main function of IL-8 is the recruitment of neutrophils in response to tissue damage caused by infection or injury (16). Whereas infection generally led to a significant increase in IL-8 production, the models infected with the piliated strains F3 Pil+ and N159 showed the highest secretion of IL-8, while the non-adherent N191 showed the lowest. The differences in IL-8 production between the models infected with F3 Pil-, F3 Pil+, or N159 were not statistically significant (Fig. 4A, B). The high secretion of IL-8 correlates in part with the observed adherence and tissue damage (Fig. 2B, D).

**Figure 4.**
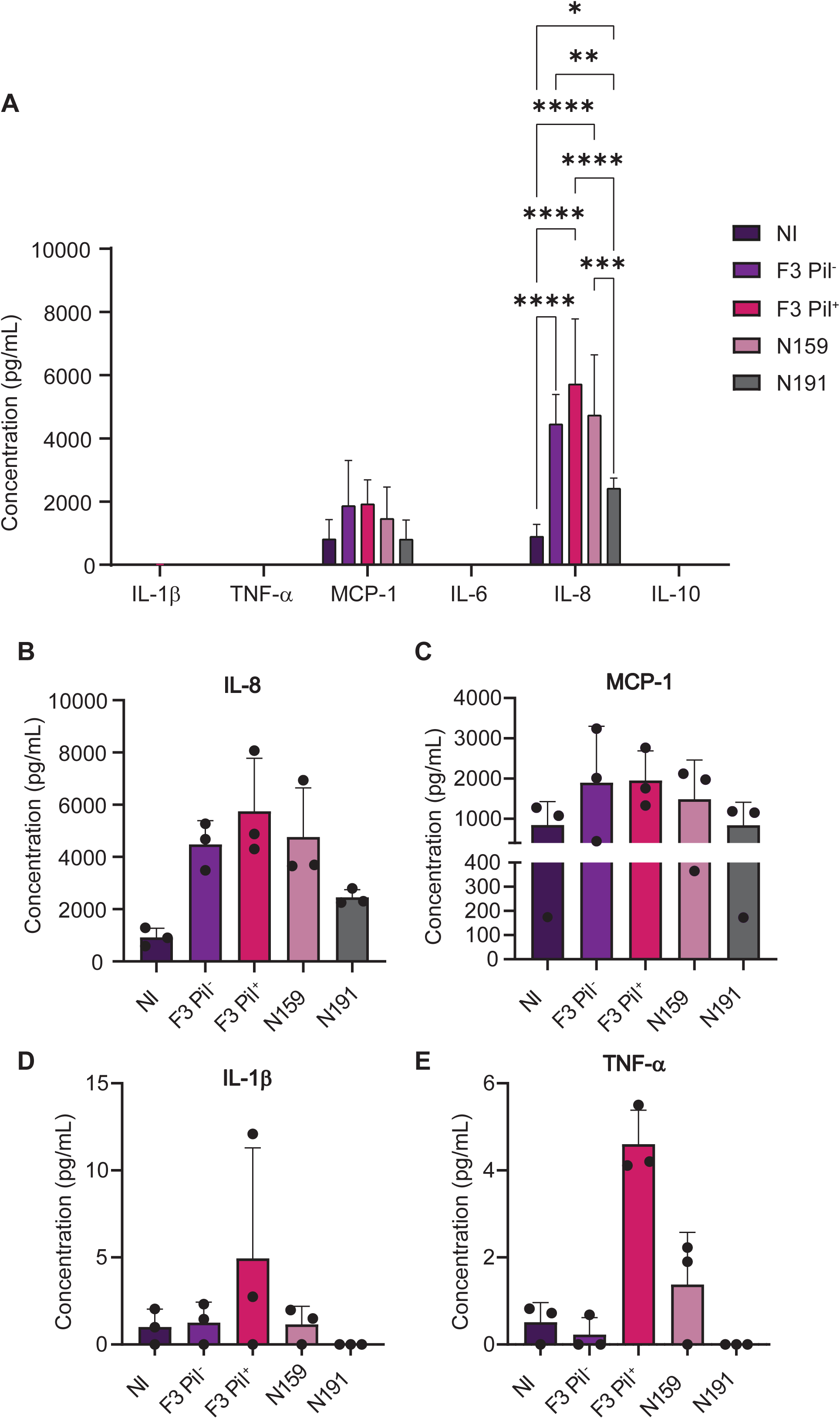
Infection of corneal epithelium models *N. gonorrhoeae* MS11 leads to an increase in IL-8 and TNF-α secretion. (A-E) Different cytokines and chemokines secreted by the corneal epithelial cells were measured in the medium collected from the basolateral side of the tissue models after 72 hours of infection using a personalized Legendplex cytokine panel to assess the secretion of interleukin (IL)-8 (B), monocyte chemotactic protein 1 (MCP-1) (C), IL-1β (D), and TNF-α (E). The graphs represent the mean values ± SD from three independent replicates. Statistical analysis was performed using 2-way ANOVA and Tukey’s multiple comparison tests. * - p≤0.05, ** - p≤0.01, *** - p≤0.001, and **** - p≤0.0001.

The secretion of MCP-1, another chemokine responsible for the recruitment of monocytes and macrophages, was also observed in all infected models, although without statistically significant differences among them and non-infected models. However, similar to the IL-8 secretion, F3 Pil-, F3 Pil+, and N159 strains showed the highest values, while N191 showed the lowest (Fig. 4C). Regarding IL-1β and TNF-α, we detected amounts less than 10 pg/ml in the infected tissue models with F3 Pil+, N159, and F3 Pil-. For the N191 strain, values were under the detection limit (Fig. 4D, E).

The infection, therefore, induces the secretion of high amounts of IL-8 that correlate with the ability of bacteria to adhere to the tissue, and are the highest for the strains that express a functional pilus.

### Trifluoperazine decreases bacterial adherence and secretion of IL-8

We next explored whether our multilayer corneal epithelium infection models are suitable for drug testing. For this, we used trifluoperazine, a small molecule that belongs to the group of phenothiazines and is currently used as an antipsychotic agent. Previously, trifluoperazine was shown to have antimicrobial properties, especially against bacteria that express the type IV pilus (17). In previous studies, trifluoperazine disrupted pilus-dependent function, such as twitching mobility, aggregation, and adherence to inert surfaces or cell surfaces in *N. meningitidis* (18). In our study, we assessed the adherence to the corneal epithelium tissue models and the IL-8 secretion after the treatment with trifluoperazine, using highly piliated N159 *N. gonorrhoeae* MS11 derivative.

In the first approach, we infected the tissue models with the N159 strain for 1 h, administered trifluoperazine (40 µM) for another hour, and then removed the medium and the drug, allowing the infection to proceed for another 72 h (post-treatment (PostTr)). In the second approach, bacteria and the drug were simultaneously added to the models for 1 h, after which they were removed by washing, followed by additional 72 h of infection (simultaneous treatment (SimTr)) (Fig. 5A).

**Figure 5.**
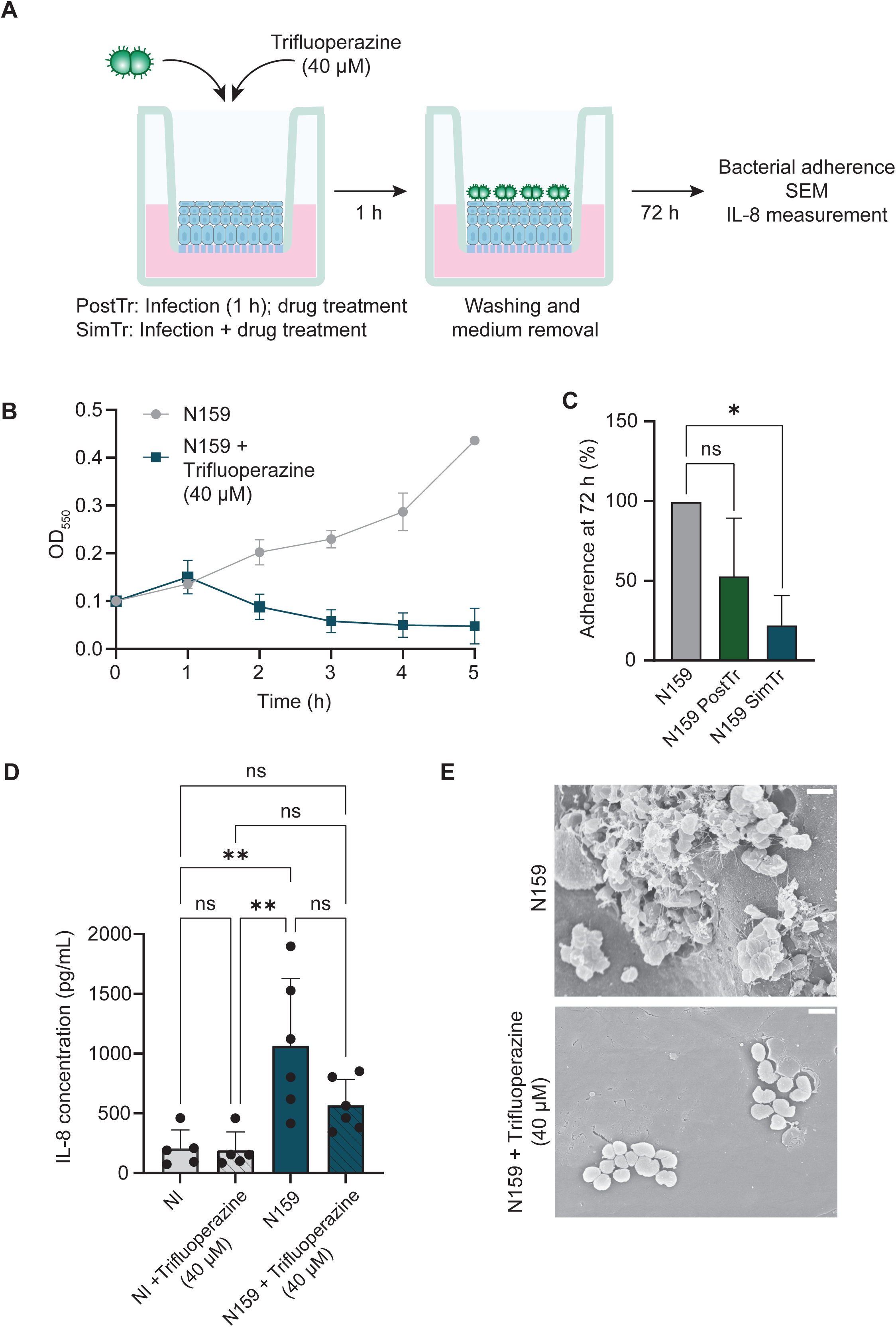
Trifluoperazine decreases bacterial adherence to the corneal epithelium models in the early stages of infection. (A) Schematic representation of two different approaches to test the effect of trifluoperazine. In the PostTr approach, models were first infected with N159 bacteria for 1 h, washed, and treated with 40 µM trifluoperazine for 1 h, followed by another wash and incubation for 72 h under airlift conditions. For the SimTr approach, the bacteria were treated with 40 µM trifluoperazine simultaneously with infection for 1 h, after which the models were washed and kept for 72 h under airlift conditions. (B) To measure the growth of N159 bacteria without and with 40 µM trifluoperazine, bacteria were grown in the PPM+ medium until they reached an OD_550_ = 0.4, diluted to an OD_550_ = 0.1, and 40 µM trifluoperazine was added to one culture. OD_550_ was measured every hour for 5 h. The graph represents mean values ± SD of at least three independent replicates. (C) Models were prepared as described in (A) and solubilized with 1 % saponin solution. The CFU/ml was calculated from the number of colonies growing after plating of serial dilutions of the suspension on GC agar plates. The results were normalized to the non-treated models. The graph represents mean values ± SD of at least three independent replicates. (D) The concentration of IL-8 was measured in the basolateral medium of the non-infected (NI) and N159-infected corneal tissue models, with and without 40 µM trifluoperazine, using ELISA. The graph shows the mean value of at least three independent replicates. Statistical analysis was performed using 2-way ANOVA and Tukey’s multiple comparison tests. * - p≤0.05, ** - p≤0.01. (E) Scanning electron microscopy images of the corneal epithelium models infected with N159 with and without the 1 h treatment with 40 µM trifluoperazine. Scale bar is 1 µm.

To control for the bactericidal effects of trifluoperazine, we monitored bacterial growth and survival in the presence of the drug. Starting from the second hour, trifluoperazine strongly affected the growth of N159 (Fig. 4B). However, analysis of bacterial survival after 1 h of treatment with different trifluoperazine concentrations in the PostTr and SimTr setups revealed interesting differences. The drug had no significant effect on bacterial survival when bacteria were first allowed to form microcolonies for 1 h (Supp. Fig. 2A). On the other hand, we observed a concentration-dependent decrease in the number of surviving bacteria when the drug was added directly to the bacterial suspension for 1 h, without previous incubation that would let bacteria aggregate (Supp. Fig. 2B). In both cases, treatment with gentamicin completely eradicated the bacteria.

Additionally, we studied the effect that trifluoperazine has on *N. gonorrhoeae* microcolonies. We incubated the suspension of N159 bacteria in a tissue culture plate for 2 h, which was sufficient for microcolony formation, and then added trifluoperazine in increasing concentrations, measuring the surface area of the microcolonies before and after 1 h of drug treatment. The drug significantly diminished the size of the microcolonies (Supp. Fig. 2C), though this did not correlate with the bactericidal effect (Supp. Fig. 2A). Gentamicin, on the other hand, did not affect bacterial microcolony size even though no bacteria survived the treatment (Supp. Fig. 2C).

In agreement with these observations, we observed no significant difference in N159 adherence to the corneal tissue models in PostTr setup, whereas the simultaneous addition of trifluoperazine significantly diminished the number of adherent bacteria (Fig. 5C). Accordingly, the drug treatment led to a reduction in IL-8 secretion in the infected corneal epithelium models (Fig. 5D). In addition, the SEM showed that, after addition of trifluoperazine, N159 microcolonies on the surface of the tissue models were reduced to only a few bacteria, with no visible pilus present (Fig. 5E).

Corneal epithelium tissue models, therefore, represent a reliable tool to test drugs that affect bacterial pilus formation and adherence. One of these drugs is trifluoperazine, which can reduce gonococcal infectivity by affecting pilus formation.

## Discussion

In this work, we show the application of a multilayer corneal epithelium tissue model for the study of the *N. gonorrhoeae* infection. These models were developed using an immortalized human corneal epithelial cell line (hTCEpi) grown on a polycarbonate membrane scaffold. hTCEpi expresses a human telomerase reverse transcriptase (hTERT), leading to the activation of telomerase, which prevents the shortening of telomeres and the telomere-dependent senescence in the cells without altering their differentiation capacity (19). Under airlift conditions, these models exhibit differentiation markers of the corneal epithelium and natural desquamation comparable to the native cornea tissue (10). However, for infection studies, additional parameters like the tissue permeability and integrity are also important. Our multilayer corneal epithelium models have a permeability of 8 % and transepithelial electrical resistance (TEER) values of 327 Ω*cm^2^ on day 7 after introducing airlift conditions, which correlates with values measured for human cornea (200 Ω*cm^2^) (20), making them suitable to be used as infection models.

One of the most important roles of the type IV pilus of *N. gonorrhoeae* is to enable bacteria to adhere to tissues and form microcolonies (21, 22). In our experiments using hTCEpi-based corneal tissue models, we observed that the adherence of the piliated F3 Pil+ and N159 strains was indeed the highest, as expected, compared to lower adherence of F3 Pil- and especially N191 (Fig. 2B). N191 strain expresses pilus and has an intact recA gene but has both PilC genes mutated or deleted. These mutations in PilC, which is the subunit located at the tip of the pilus and described as the adhesin that allows the contact of the bacteria with the target tissue and other bacteria, are responsible for the reduced adherence capacity of this strain, even though PilE is expressed (Fig. 2A) (6). In our infection model, we could faithfully reproduce the effect of the PilC defect and consistently observe low adherence of N191 bacteria.

The differences in adhesion between piliated and non-piliated/non-adherent bacteria were particularly noticeable at the earliest time point of 2 h. At later time points, the number of adherent bacteria increased for all strains (Fig. 2B), probably due to the multiplication of already-attached GC (Fig. 3A). Interestingly, F3 Pil- showed better adherence than the N191 strain, indicating that the presence of a non-functional pilus might interfere with the tissue contact. Both strains still interacted with the tissue, however, possibly through opa proteins (23), since we did not specifically select opa-negative bacterial colonies for infection (Fig. 2A). An earlier report showed that non-piliated bacteria did not adhere to human corneal explants, and that the adherence was independent of the opa status (8). However, in that study, bacteria were incubated with the tissue for 1 h, whereas in our experiments, the incubation time was 2 h. It is possible that longer incubation allowed bacteria to interact with the tissue using opa proteins, and that the differences in adherence between piliated and non-piliated bacteria would be more apparent at shorter incubation times.

The same earlier study reported thinning and exfoliation of the human cornea explants upon infection (8). *N. gonorrhoeae* was also shown to disrupt the apical junctions of polarized epithelial cells (24, 25) and induce epithelial cell exfoliation in cervical tissue explants (26). Contrary to this, we did not see major differences in tissue permeability changes over time between infected and non-infected hTCEpi cornea models that would be indicative of tissue thinning or exfoliation (Fig. 2C). The cytotoxicity also did not seem to strictly depend on the presence of the pilus but mostly correlated with the number of adherent bacteria (Fig. 2D). This might be due to the differences between tissue explants and models, with models showing greater robustness in infection scenario.

The formation of microcolonies is important for the survival of GC in the host because it facilitates invasion and reduces the effectiveness of antibiotics, as they are less able to diffuse within the microcolony. In our study, we observed tissue damage and formation of pits under the microcolonies formed by highly piliated N159. This is reminiscent of the observations made using human cornea explants (8). Tissue damage caused by GC attachment could be related to cornea perforation, one of the most severe complications in patients with untreated eye infections caused by *N. gonorrhoeae* (27, 28). Likewise, SEM images at 72 h revealed that F3 Pil- forms occasional biofilm-like patches on the surface of the models. No pilus was visible, but there was an apparent tissue destruction beneath these patches (Fig. 3B), which could explain the high cytotoxicity measured for models infected with this strain (Fig. 2D). Interestingly, however, the lowest percentage of cytotoxicity was seen after F3 Pil+ infection and not after N191 infection, suggesting that the cell death caused by GC in the corneal epithelium tissue is not exclusively pilus-dependent, and the role of other virulence factors is also relevant.

We also measured the concentration of different cytokines and chemokines secreted upon infection by the corneal epithelial cells. The pattern of cytokines depends on the tissue in question. In T-cells, the pilus-dependent adherence of *N. gonorrhoeae* induces the activation and proliferation of these cells and the production of anti-inflammatory IL-10 (29). In mature macrophages incubated with piliated GC, the infection causes an increased secretion of the pro-inflammatory cytokines IL-6, TNF-α, GRO-α, MIP-1α, and RANTES, with no effect on IL-8 (30). In the male urethra (31), there is an increased expression of IL-8, IL-6, and TNF-α in urine before the start of symptoms after GC infection, and in fallopian tubes, IL-1α, IL-1β, and TNF-α are upregulated (32). Our previous work using 3D models that mimicked endometrial and urethral tissue showed that urethral tissue models secreted high amounts of IL-8, IL-6, and TNF-α, whereas endometrial tissue models secreted significantly less of the same cytokines (11). There is little known, however, about the cytokine secretion of corneal epithelium upon GC infection. We observed an increase in IL-8 and TNF-α secretion in infected cornea models, which correlated with the number of adherent bacteria. MCP-1 secretion was also increased, but this did not seem to be related to the infection. Only small amounts of IL-1β, and no IL-6 or IL-10, were produced upon infection (Fig. 4). These results point to a specific, pro-inflammatory response of corneal tissue to GC infection, mostly directed at the recruitment of neutrophils, which corresponds to the clinical picture of the disease.

Finally, we tested the effects of trifluoperazine on pilus-mediated adherence and infection using corneal tissue models. Trifluoperazine reduces bacterial piliation by affecting the establishment of the sodium gradient through alteration of the activity of the Na^+^-pumping NADH-ubiquinone oxidoreductase complex (18). This complex creates a Na^+^ gradient across the cell membrane using energy released by the oxidation of NADH and the reduction of quinone (33). Trifluoperazine was described to induce pilus retraction and dispersal of microcolonies in *N. meningitidis* (18). We demonstrated a similar effect on *N. gonorrhoeae* in liquid culture (Supp. Fig. 2). However, in tissue models, we observed that when we allowed piliated GC to adhere and form microcolonies on the tissue models for 1 h before the addition of trifluoperazine, the drug was not as efficient as when added simultaneously with bacteria (Fig. 5C). This is in agreement with the role of the pilus only in the initial attachment to the tissue, with adherence to host cells being mediated by other adhesins as the infection progresses.

Our study demonstrates for the first time the usefulness of hTCEpi-based tissue models for infection research, shedding light on several important aspects of the infection of the cornea with *N. gonorrhoeae* and the role of type IV pilus in this process. In comparison to the primary tissue in the form of explants, corneal epithelial tissue models are more accessible and show lower variability, while still faithfully mimicking the original tissue, making them suitable for high-throughput research and drug screening.

## Acknowledgments

We thank Leonie Speth for technical assistance. This study was supported by the Deutsche Forschungsgemeinschaft (DFG, German Research Society) through a GRK2157/1+2 grant to VK-P, which included the salary of LK. The funders had no role in study design, data collection and analysis, decision to publish, or preparation of the manuscript.

## Data Availability

The sequencing data related to this study is available and can be accessed from BioProject: PRJNA1356842.

## Supplementary figure legends

**Supplementary Figure 1.** Pilin protein in F3 Pil- strain has several deletions. Amino acid sequences were translated using ExPasy according to the nucleotide sequences obtained from whole genome sequencing and aligned to the reference MS11 sequence (NCBI (CP003909)) using Clustal Omega.

**Supplementary Figure 2.** Trifluoperazine has a bactericidal effect as well as causing disaggregation of GC microcolonies. (A-B) The same number of N159 bacteria used for the infection of corneal epithelium models (6x10^6^) was treated with increasing concentrations of trifluoperazine. For the PostTr approach, trifluoperazine was added after 1 h of incubation (37 °C, 5 % CO_2_) for 1 h. For the SimTr approach, the bacteria were treated simultaneously with infection for 1 h. Serial dilutions were plated to calculate the CFU/ml. 150 µg/ml of gentamicin was used as a positive control. (C) N159 bacteria were grown in PPM+ medium until they reached an OD_550_ of 0.4, diluted to an OD_550_ of 0.1 in DMEM F12, transferred into a 24-well plate, and incubated for 2 h (37 °C, 5 % CO_2_). After this time, the bacteria were treated with increasing concentrations of trifluoperazine or with gentamicin for 1 h. Three images of each of the triplicates were made before and after the treatment using a phase-contrast microscope, and the area index of GC microcolonies was calculated using FIJI. The graph represents the mean values ±SD of three independent replicates. Statistical analysis was performed using 2-way ANOVA and Tukey’s multiple comparison tests. * - p≤0.05, ** - p≤0.01, and **** - p≤0.0001.

