## Supplemental Figures for "The role of type IV pilus in the interaction of *Neisseria gonorrhoeae* with a corneal epithelium tissue model"

### Supplementary Figure 1

```
01821_PilE      MNTLQKGFTLIELMIVIAIVGILAAVALPAYQDYTARAQVSEAILLAEGQKSAVTEYYLN 60
F3_Pil-         ----------------------------------------------------PPARKFPK 8
F3_Pil+         ----------------------------------------------------PPARKFPK 8
N159            ----------------------------------------------------PPARKFPK 8
N191            ----------------------------------------------------PPARKFPK 8
                                                                      :. : :

01821_PilE      HGKWPENNT------------------SAGVASPPTDIKGKYVKEVEVKNGVVTATMLSS 102
F3_Pil-         PSFWPKVKNQPLPSIT-ITANGRKTTLLPAWHPPPTKSKANMFRKLKSQKASLPPKWLQP 67
F3_Pil+         PSFWPKVKNQPSPSIT-ITAYGRKTTALPAWHPPPT-SKANMLSKLRSQTASLPPK-NQT 65
N159            PSFWPKVKNQPSPSIT-ITAYGRKTTALPAWHPPPT-SKANMLSKLRSQTASLPPK-NQT 65
N191            PSFWPKVKNQPSPSIT-ITAYGRKTTTLPAWHPPPT-SKANMLSKLRSQTASLPPKWLQP 66
                 . **: :.       *            .   ***  *.: . ::. :.. :  .  .

01821_PilE      GVNNEIKGKKLSLWGRRENGSVKWFCGQPVTRADDDTVA---------DAKDGKEIDTKH 153
F3_Pil-         A-TKKSKTKNSP-------------CGPSVKTVR-NGSADSRLRAPTTTPLPTPKTAKKS 112
F3_Pil+         A-TKKSKTKKLSLWAKRQDGSVKWFCGQPVTRN---AKADDTVT---KAGNDNEKINTKH 118
N159            A-TKKSKTKKLSLWARREAGSVKWFCGQPVTRN---DKANVTDD---ADVTGNDKIETKH 118
N191            A-TKKSKTKKLSLWAKRQDGSVKWFCGQPVTRN---AKADDTVT---KAGNDNEKINTKH 119
                . .:: * *:               **  *.       *               :  .*

01821_PilE      LPSTCRDKASDAK--- 166
F3_Pil-         TPSTCRQPAAITLMPN 128
F3_Pil+         LPSTCRDNFDAS---- 130
N159            LPSTCRDNFDAS---- 130
N191            LPSTCRDNFDAS---- 131
                 *****:    :
```

### Supplementary Figure 2

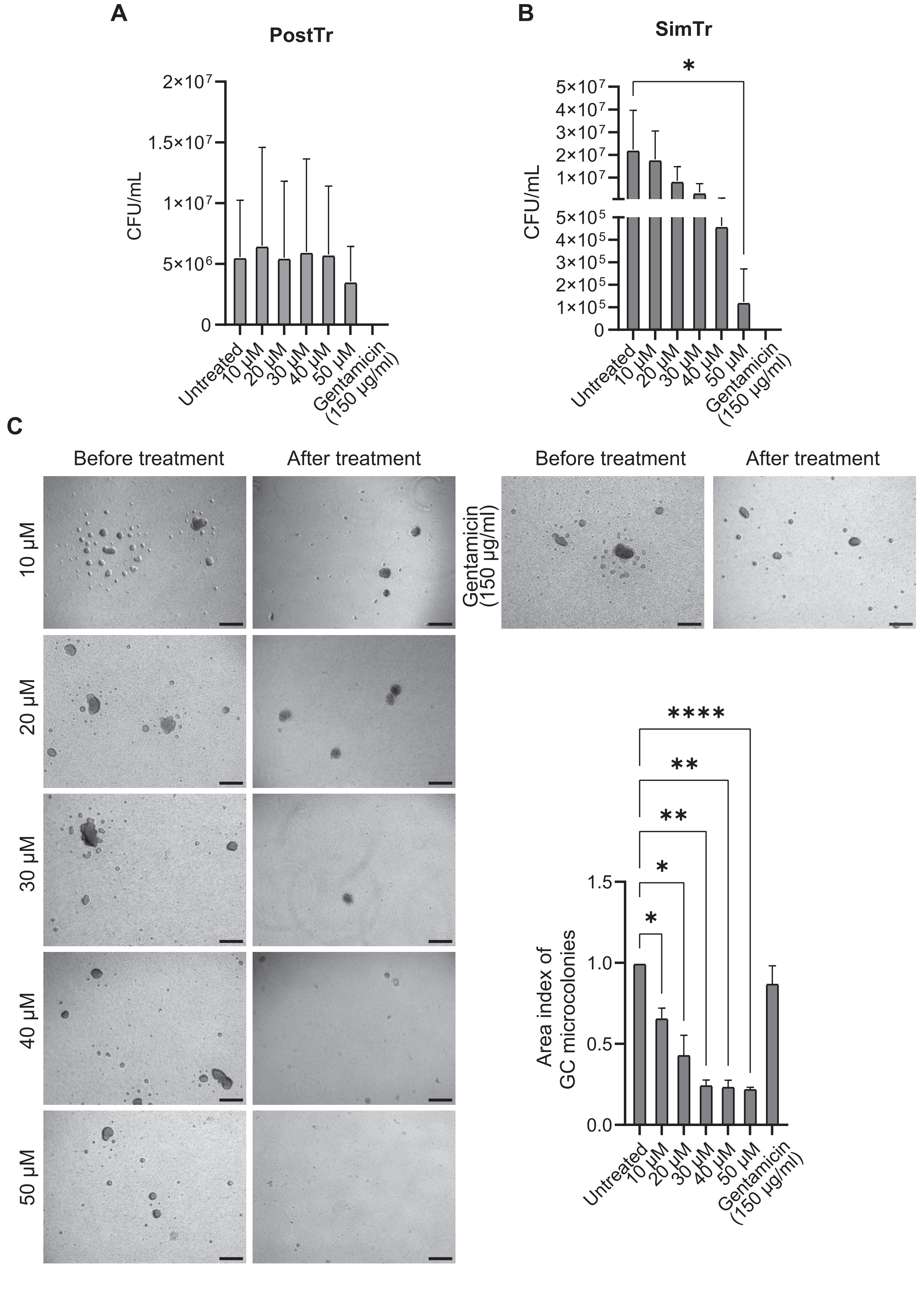

### Supplementary figure legends

**Supplementary Figure 1.** Pilin protein in F3 Pil- strain has several deletions. Amino acid sequences were translated using ExPasy according to the nucleotide sequences obtained from whole genome sequencing and aligned to the reference MS11 sequence (NCBI (CP003909)) using Clustal Omega.

**Supplementary Figure 2.** Trifluoperazine has a bactericidal effect as well as causing disaggregation of GC microcolonies. (A-B) The same number of N159 bacteria used for the infection of corneal epithelium models ($6x10^6$) was treated with increasing concentrations of trifluoperazine. For the PostTr approach, trifluoperazine was added after 1 h of incubation (37 °C, 5 % $CO_2$) for 1 h. For the SimTr approach, the bacteria were treated simultaneously with infection for 1 h. Serial dilutions were plated to calculate the CFU/ml. 150 µg/ml of gentamicin was used as a positive control. (C) N159 bacteria were grown in PPM+ medium until they reached an $OD_{550}$ of 0.4, diluted to an $OD_{550}$ of 0.1 in DMEM F12, transferred into a 24-well plate, and incubated for 2 h (37 °C, 5 % $CO_2$). After this time, the bacteria were treated with increasing concentrations of trifluoperazine or with gentamicin for 1 h. Three images of each of the triplicates were made before and after the treatment using a phase-contrast microscope, and the area index of GC microcolonies was calculated using FIJI. The graph represents the mean values ±SD of three independent replicates. Statistical analysis was performed using 2-way ANOVA and Tukey's multiple comparison tests. * - $p \leq 0.05$, ** - $p \leq 0.01$, and **** - $p \leq 0.0001$.
